## Supplementary Figures for "Deficiency of Orexin Receptor Type 1 in Dopaminergic Neurons Increases Novelty-Induced Locomotion and Exploration"

### Figure 1—figure supplement 1

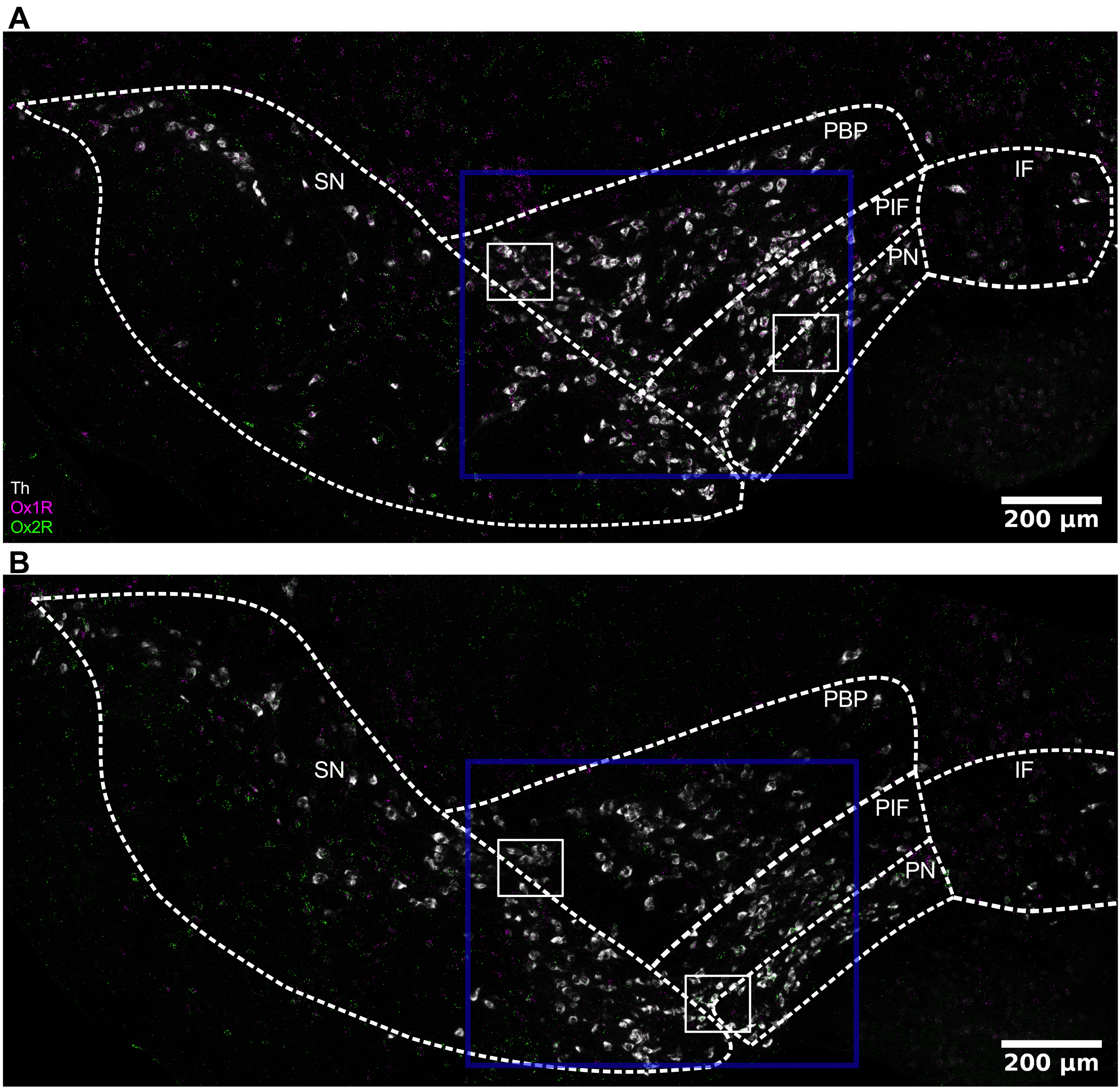


**An overview of orexin receptor expression in the substantia nigra (SN) and the ventral tegmental area (VTA).** Representative images of RNAscope *in situ* hybridization in SN and interfascicular (IF), paranigral (PN), parainterfascicular (PIF) and parabrachial (PBP) subnuclei of VTA, of **(A)** a control and **(B)** an Ox1R$^{\Delta DAT}$ mouse. The amplications of areas marked in blue squares are shown in Figure 1A for the control mouse and in Figure 1B for the Ox1R$^{\Delta DAT}$ mouse. White, tyrosine hydroxylase (Th); magenta, Ox1R; green, Ox2R. Scale bar: 200 µm. Control, n = 3; Ox1R$^{\Delta DAT}$, n = 4.

### Figure 1—figure supplement 2

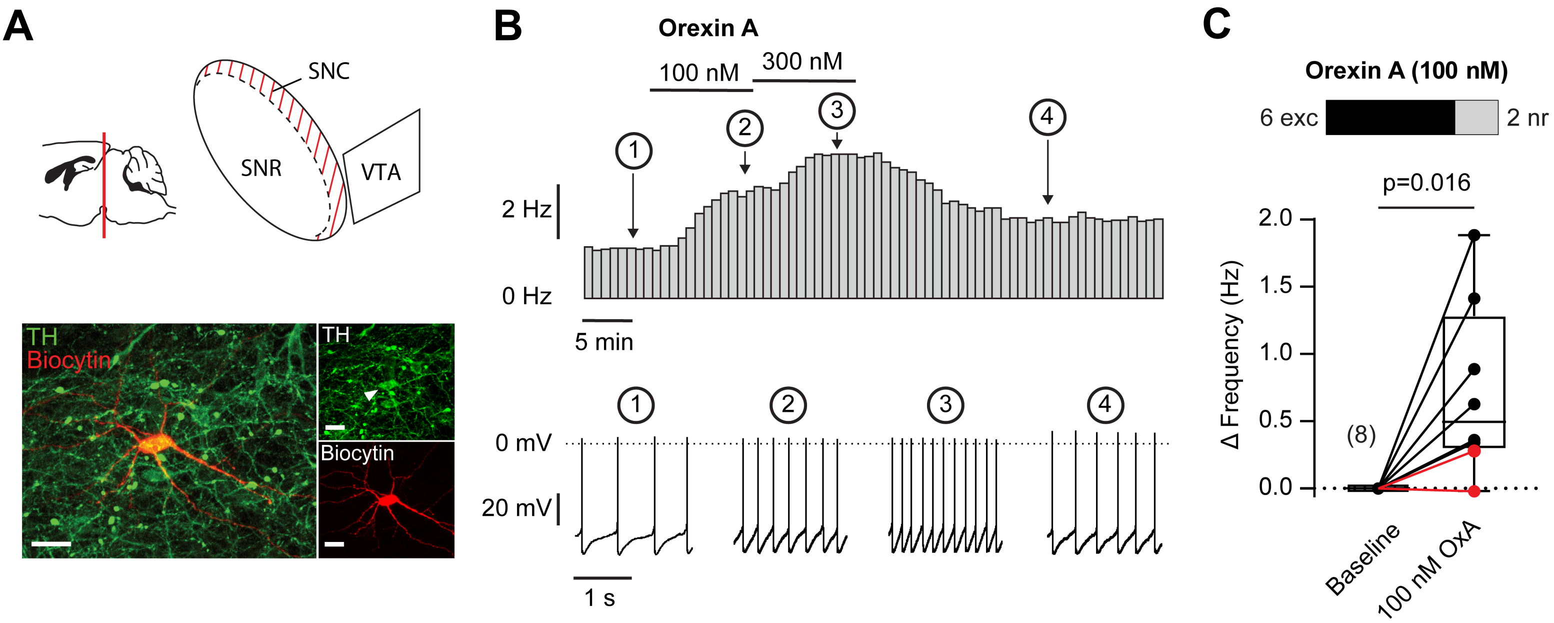


**Orexin A increases the action potential frequency of dopaminerg neurons in the substantia nigra (SN).**
Recordings were performed from dopaminergic SN neurons in acute brain slices of control mice in the perforated patch clamp configuration to preserve intracellular signaling pathways. In all recordings, GABAergic and glutamatergic synaptic input was pharmacologically blocked (see Materials and Methods). Orexin A was bath applied at 100 nM and 300 nM for 8 min for each concentration. (**A**) Top: Schematic illustration of the substantia nigra where the recordings have been performed (red crosshatched). Bottom: Image of a recorded dopaminergic SN neuron, which was filled with biocytin after converting the perforated patch clamp configuration to the whole-cell configuration. The neuron was double-labeled with biocytin-streptavidin (red) and an anti-serum against tyrosine hydroxylase (green). Scale bar: 20 µm. **(B)** Orexin A effect on action potential firing rate of dopaminergic SN neurons. Rate histograms (bin width: 50 s) (top) and representative sections of the original recordings (bottom). The numbers mark the recording sections displayed in high time resolution as original traces (bottom). (**C**) Top: The stacked bars show the number of individual neurons in which the increase in action potential frequency was three times larger than the standard deviation (3 σ criterion ≙ Z-score of 3) of the control, thus defining them as responsive (see Materials and Methods) (exc, excited; nr, not responsive). Bottom: The population effect of orexin A on the action potential frequency was measured as the orexin A induced change in frequency (Δ frequency) and tested for significance using a one sample Wilcoxon matched-pairs signed rank test ($p = 0.016$, $n = 8$). The horizontal lines in the box plots show the median. The whiskers were calculated using the Tukey method. Recordings not responding to 100 nM orexin A are indicated in red. *n* values are given in brackets. SNR, substantia nigra pars reticulata; SNC, substantia nigra pars compacta; VTA, venral tegmental area; OxA, orexin A.

### Figure 1—figure supplement 3

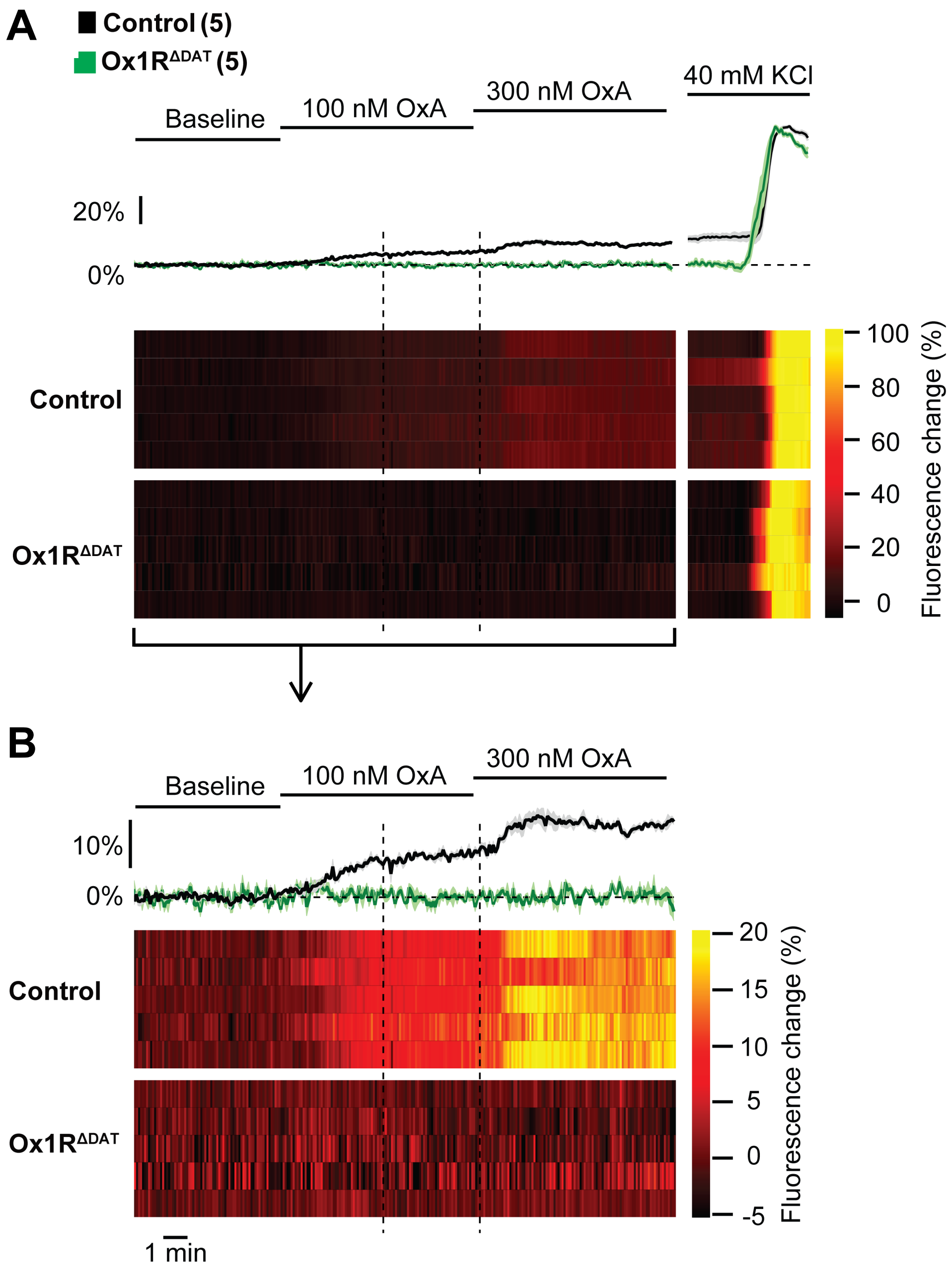


**Orexin A and high $K^+$ saline effect on $[Ca^{2+}]_i$ of dopaminergic neurons in the substantia nigra (SN) of control and Ox1R$^{\Delta DAT}$ mice which express GCaMP6s.**

Change in GCaMP6 fluorescence induced by orexin A (100 nM and 300 nM) and high (40 mM) $K^+$ saline, to which the orexin A responses were normalized for quantification (see Figure 1L). As described in Materials and Methods, the orexin A response can be expressed for each neuron individually as a percentage of its high $K^+$-induced GCaMP6 fluorescence. This value is a solid reference point reflecting the GCaMP6 fluorescence at maximal voltage-activated $Ca^{2+}$ influx. The $Ca^{2+}$ concentration at this point is exceptionally high and not typically reached under physiological conditions. Therefore, as shown in (A), the physiologically relevant responses may, at first glance, appear to be relatively small when plotted together with the high $K^+$ response. In **(A)** and **(B)**, for completeness, different presentation options and scales are exemplified using the 5 orexin A-responsive dopaminergic SN neurons from control mice and the 5 not orexin A-responsive dopaminergic neurons of Ox1R$^{\Delta DAT}$ mice neurons from Figure 1K (Heat maps of all recorded neurons are shown in Figure 1—figure supplement 4). Top: mean responses. Error bars are not displayed for clarity reasons. Bottom: Heat maps showing the Orexin A effect of the individual neurons. On the single-cell level, neurons were considered orexin A-responsive when the change in $[Ca^{2+}]_i$ was larger than three times the standard deviation (3 σ criterion ≙ Z-score of 3) of the baseline fluorescence (see Materials and Methods). OxA, orexin A.

### Figure 1—figure supplement 4

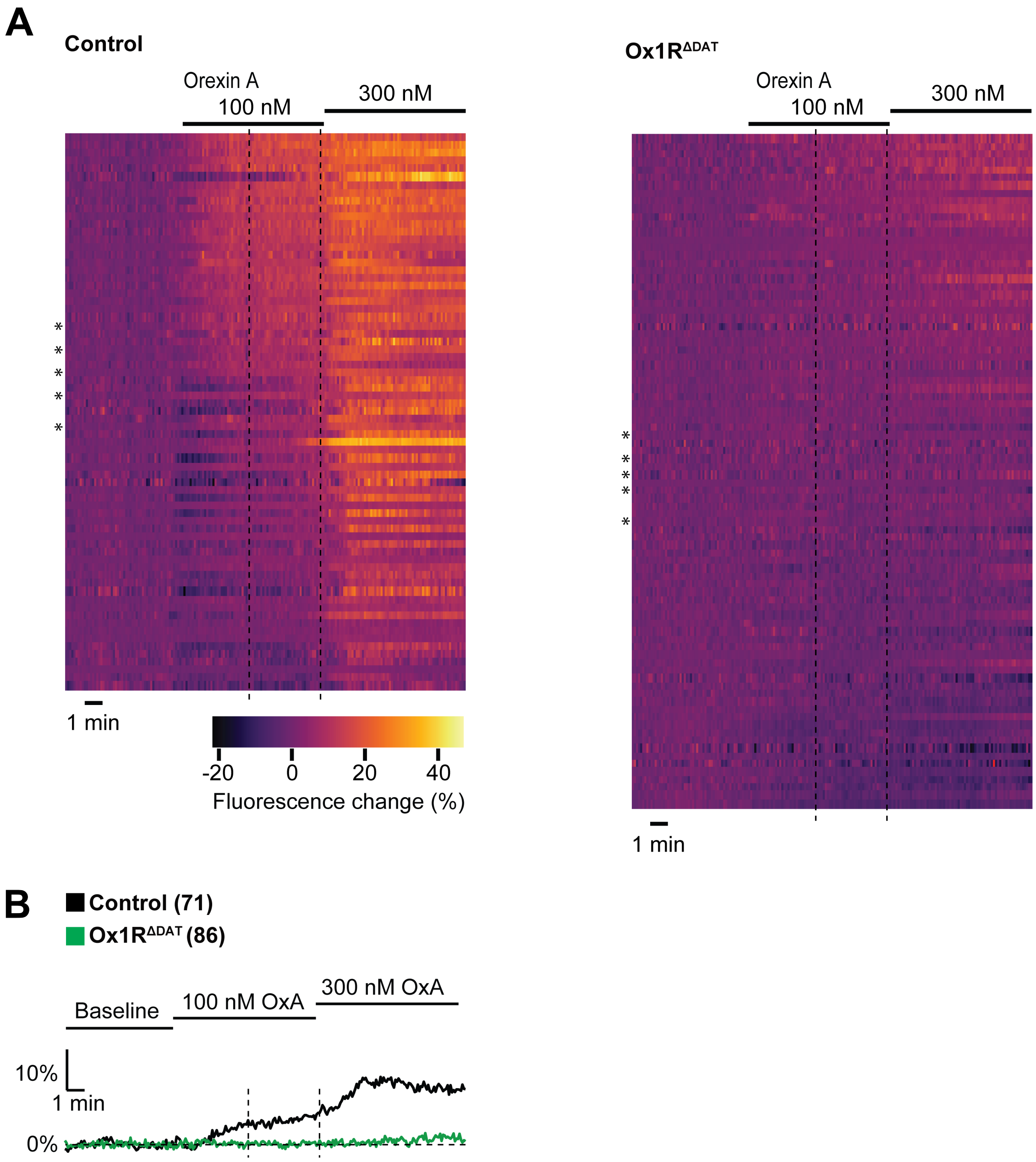


**Orexin A and high $K^+$ saline effect on $[Ca^{2+}]_i$ of dopaminergic neurons in the substantia nigra (SN) of control and Ox1R$^{\Delta DAT}$ mice which express GCaMP6s.**

**(A)** Individual heat maps showing the change in GCaMP6s fluorescence induced by 100 nM and 300 nM orexin A from dopaminergic SN neurons from (left) control (n = 71) and (right) Ox1R$^{\Delta DAT}$ mice (n = 86). The asterisks indicate the example responses shown in Figure 1K and Figure 1—figure supplement 3. **(B)** Mean responses of all recorded dopaminergic neurons of control and Ox1R$^{\Delta DAT}$ mice shown in (A). Error bars are not displayed for clarity reasons.

### Figure 2–figure supplement 1

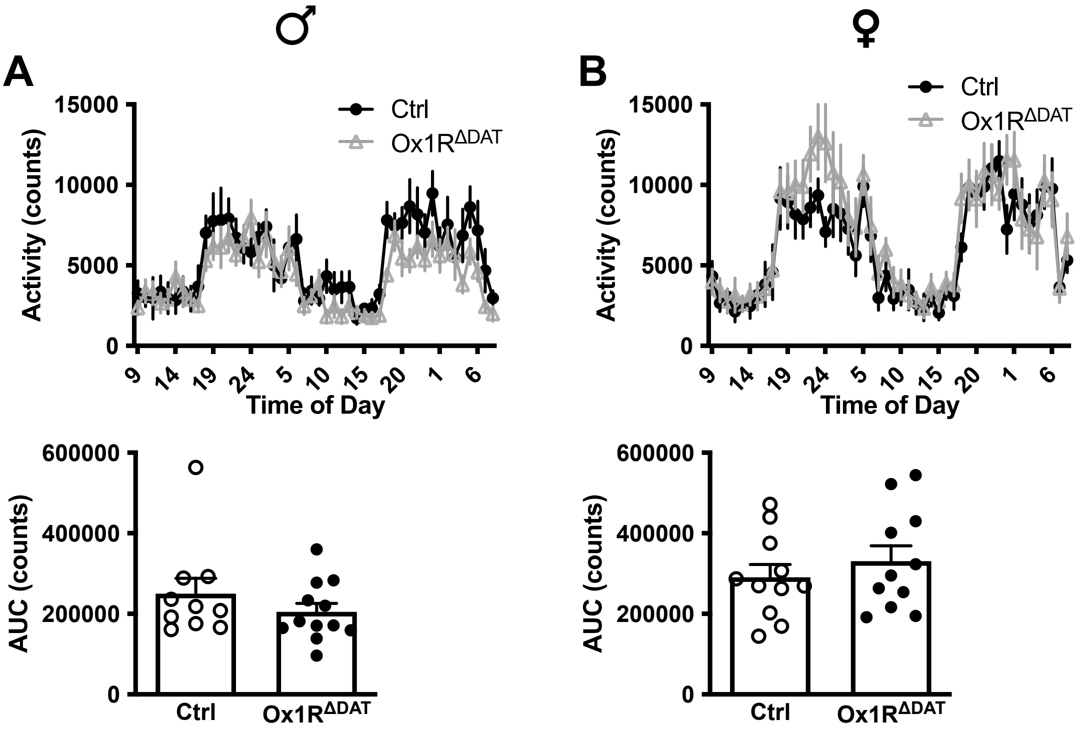


**Unaltered mouse spontanous activity in home cages by deficiency of Ox1R in dopaminergic neurons.**

Spontanous activity in home cages of **(A)** male and **(B)** female mice. Top: original counts, bottom: area under curve (AUC) was calculated for unpaired two-tailed Student's t-test. There were no significant changes between groups. Male-control, n = 10; male-Ox1R$^{\Delta DAT}$, n = 12; female-control, n = 11; female-Ox1R$^{\Delta DAT}$, n = 11. Data are represented as means ± SEM. The full statistics are provided alongside the source data.

### Figure 2–figure supplement 2

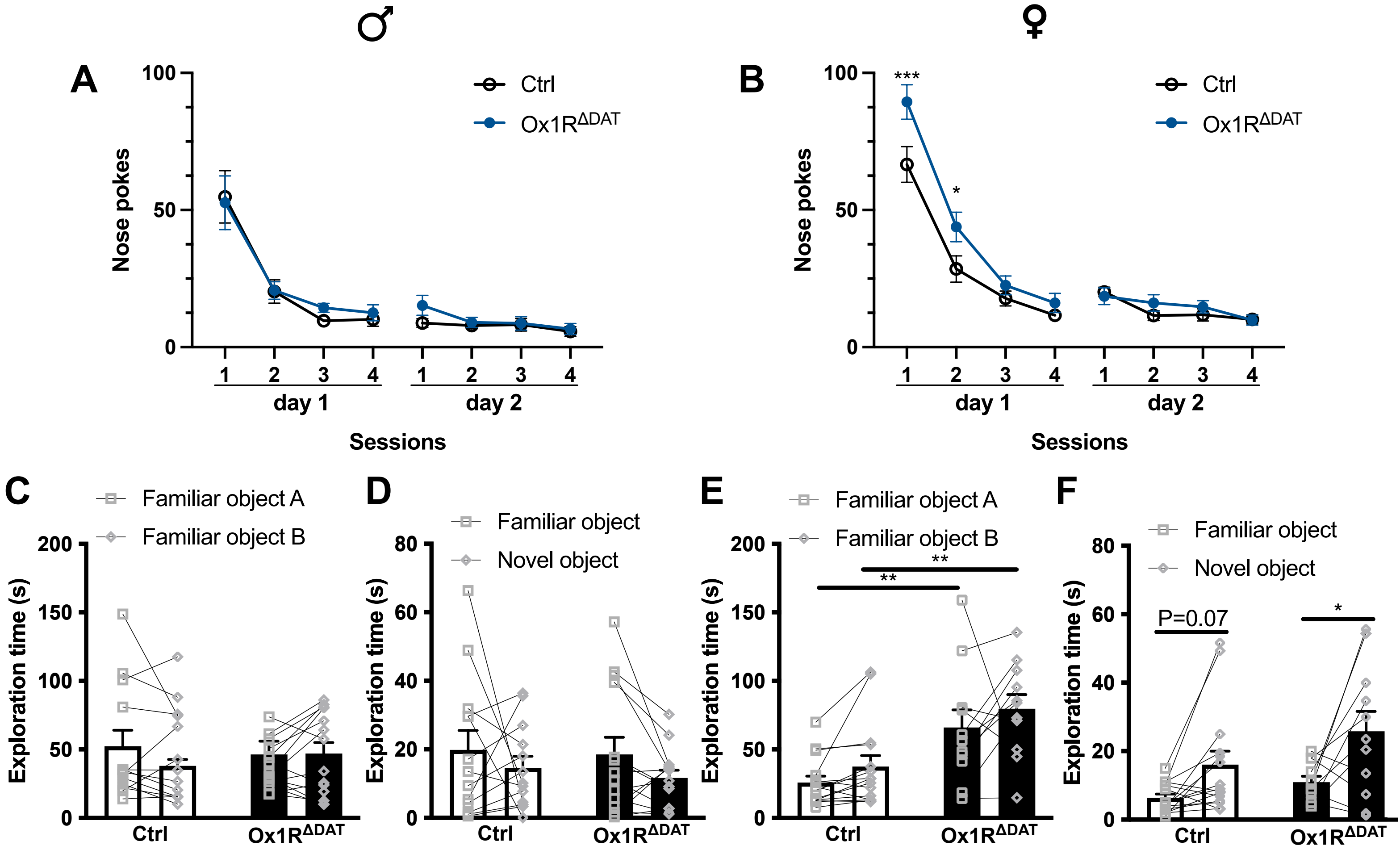


**Increased exploratory behaviours of female Ox1R$^{ΔDAT}$ in the hole-board (HB) test and the novel object test (NOT).** **(A)** Nose pokes of male mice in the hole-board test. Control, n = 10; Ox1R$^{ΔDAT}$, n = 9. **(B)** Nose pokes of female mice in the hole-board test. Control, n = 8; Ox1R$^{ΔDAT}$, n = 7. Exploration time of the objects by male mice at **(C)** acclimation phase and **(D)** novelty-exposure phase in the novel object test. Control, n = 13; Ox1R$^{ΔDAT}$, n = 14. Exploration time of the objects by female mice at **(E)** acclimation phase and **(F)** novelty-exposure phase in the novel object test. Control, n = 15; Ox1R$^{ΔDAT}$, n = 11. Data are represented as means ± SEM. *p < 0.05; **p < 0.01; ***p < 0.001; as determined by two-way ANOVA followed by Sidak's post hoc test. (A) Mixed-effect analysis (matching across time) was used because two-way ANOVA could not be performed due to one missing value in the Ox1R$^{ΔDAT}$ group in the last session: time, p < 0.0001; genotype, p = 0.53; time x genotype, p = 0.98. Two-way ANOVA (matching across time or object): (B) time: F (7, 91) = 92.57, p < 0.0001; genotype: F (1, 13) = 7.41, p = 0. 02; time x genotype: F (7, 91) = 3.07, p = 0.006; subject: F (13, 91) = 2.15, p = 0.02; (C) object: F (1, 25) = 0.10, p = 0.76; genotype: F (1, 25) = 0.04, p = 0.55; object x genotype: F (1, 25) = 2.50, p = 0.13; subject: F (25, 25) = 5.94, p < 0.0001; (D) object: F (1, 25) = 2.45, p = 0.13; genotype: F (1, 25) = 0.23, p = 0.64; object x genotype: F (1, 25) = 0.04, p = 0.84; subject: F (25, 25) = 1.45, p = 0.18; (E) object: F (1, 24) = 2.98, p = 0.10; genotype: F (1, 24) = 16.23, p = 0.0005; object x genotype: F (1, 24) = 0.02, p = 0.89; subject: F (24, 24) = 1.94, p = 0.06; (F) object: F (1, 24) = 13.49, p = 0.001; genotype: F (1, 24) = 3.67, p = 0.07; object x genotype: F (1, 24) = 0.60, p = 0.45; subject: F (24, 24) = 1.25, p = 0.29. The full statistics are provided alongside the source data.

### Figure 2–figure supplement 3

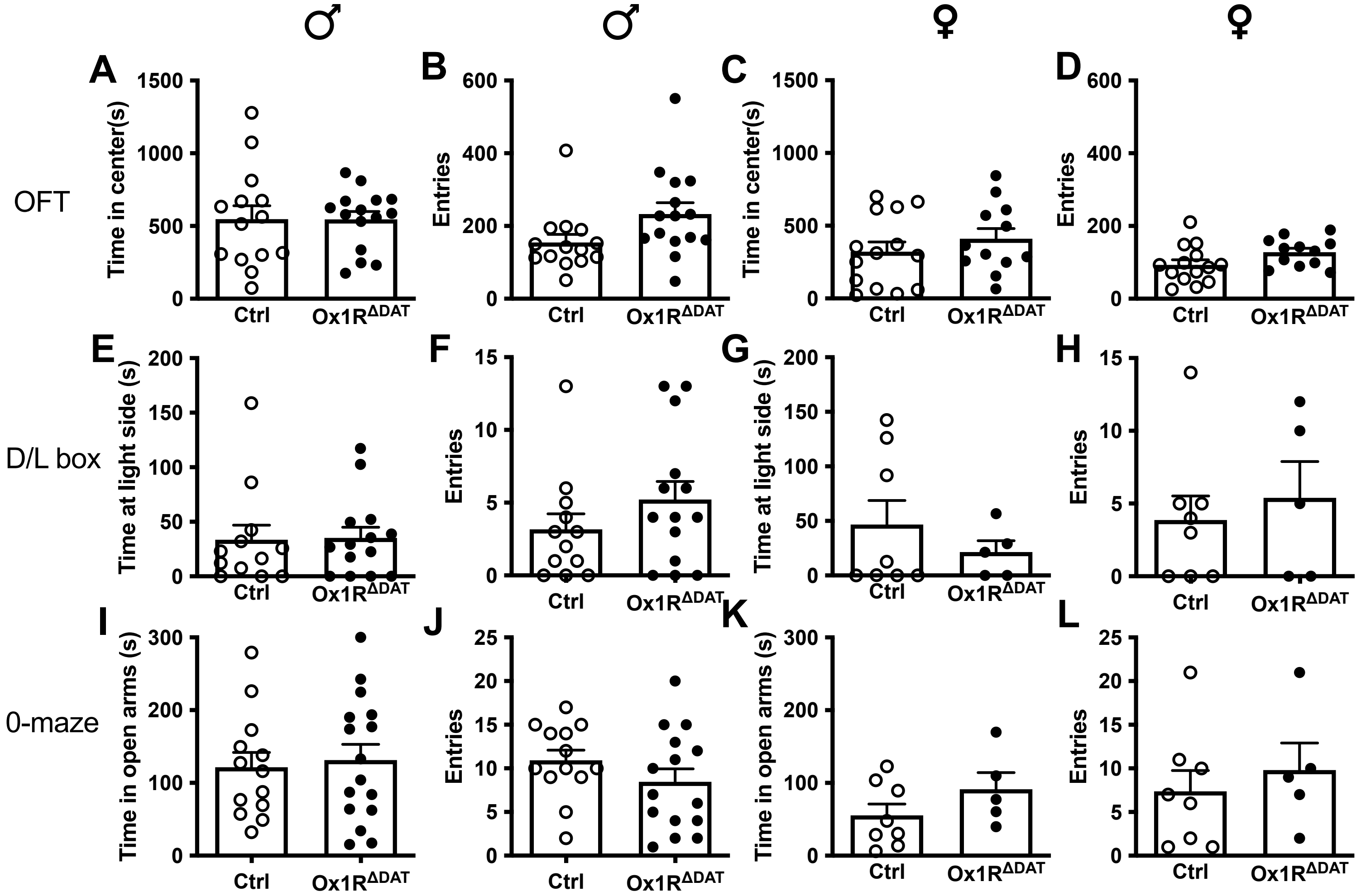


**Unaltered anxiety-related behaviours in Ox1R$^{\Delta DAT}$ mice.**

**(A)** Time spent in the center area and **(B)** times of entries into the center area by male mice in the open field test (OFT). Control, n = 14; Ox1R$^{\Delta DAT}$, n = 15. **(C)** Time spent in the center area and **(D)** times of entries into the center area by female mice in the open field test. Control, n = 14; Ox1R$^{\Delta DAT}$, n = 12. **(E)** Time spent at the light side and **(F)** times of entries into the light side by male mice in the dark/light (D/L) box test. Control, n = 12; Ox1R$^{\Delta DAT}$, n = 14. **(G)** Time spent at the light side and **(H)** times of entries into the light side by female mice in the dark/light box test. Control, n = 8; Ox1R$^{\Delta DAT}$, n = 5. **(I)** Time spent in the open arms and **(J)** times of entries into the open arms by male mice in the 0-maze test. Control, n = 13; Ox1R$^{\Delta DAT}$, n = 16. **(K)** Time spent in the open arms and **(L)** times of entries into the open arms by female mice in the 0-maze test. Control, n = 8; Ox1R$^{\Delta DAT}$, n = 5. Data are represented as means ± SEM. No significant differences between groups were detected by unpaired two-tailed Student's t-test. The full statistics are provided alongside the source data.

### Figure 2–figure supplement 4

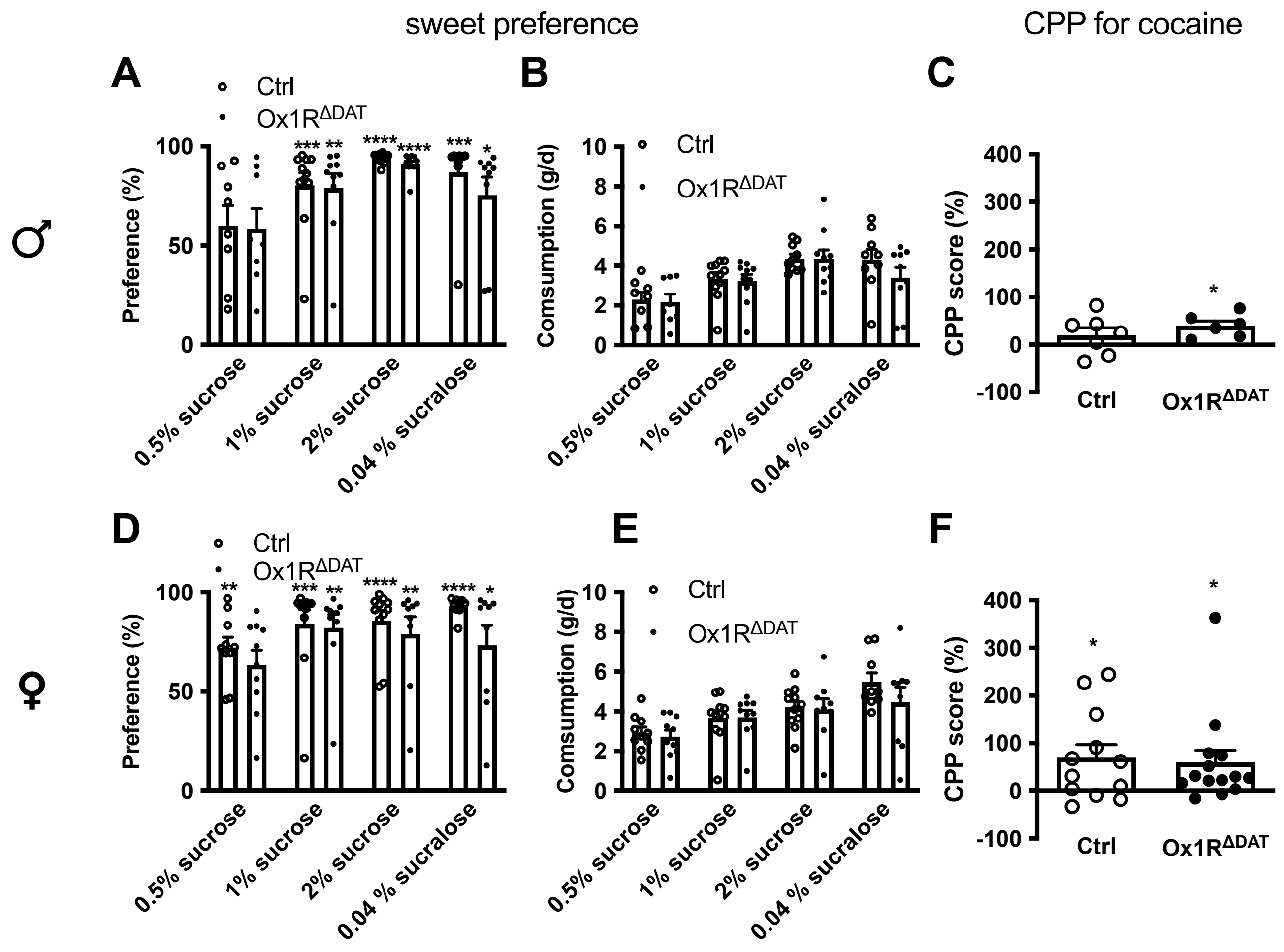


**Unaltered reward-related behaviours in Ox1R$^{\Delta DAT}$ mice.**

Sucrose/sucralose solution preference of male mice expressed as **(A)** percentages in total daily intake and **(B)** absolute values of sucrose/sucralose solution consumption in the sucrose/sucralose preference test. Control, n = 8-12; Ox1R$^{\Delta DAT}$, n = 8-10. **(C)** Conditional place preference (CPP) score of male mice in the CPP test for cocaine. Control, n = 7; Ox1R$^{\Delta DAT}$, n = 6. Sucrose/sucralose solution preference of female mice expressed as **(D)** percentages in total daily intake and **(E)** absolute values of sucrose/sucralose solution consumption in the sucrose/sucralose preference test. Control, n = 9-12; Ox1R$^{\Delta DAT}$, n = 9-10. **(F)** CPP score of female mice in the CPP test for cocaine. Control, n = 12; Ox1R$^{\Delta DAT}$, n = 14. Data are represented as means ± SEM. *p < 0.05; **p < 0.01; ***p < 0.001; ****p < 0.0001; as determined by one-sample t-test (theoretical value: A and D, 50; C and F, 0). Two-way ANOVA followed by Sidak's post hoc test: (A) Sweet solution: F (3, 67) = 7.29, p = 0.0003; genotype: F (1, 67) = 0.78, p = 0.38; sweet solution x genotype: F (3, 67) = 0.23, p = 0.87; (B) Sweet solution: F (3, 67) = 9.99, p < 0.0001; genotype: F (1, 67) = 1.05, p = 0.31; sweet solution x genotype: F (3, 67) = 0.55, p = 0.65; (D) Sweet solution: F (3, 67) = 2.46, p = 0.07; genotype: F (1, 67) = 3.65, p = 0.06; sweet solution x genotype: F (3, 67) = 0.58, p = 0.63; (E) Sweet solution: F (3, 67) = 8.37, p < 0.0001; genotype: F (1, 67) = 1.14, p = 0.29; sweet solution x genotype: F (3, 67) = 0.58, p = 0.63. Unpaired two-tailed Student's t-test: (C) p = 0.32, (F) p = 0.79. The full statistics are provided alongside the source data.

### Figure 2–figure supplement 5

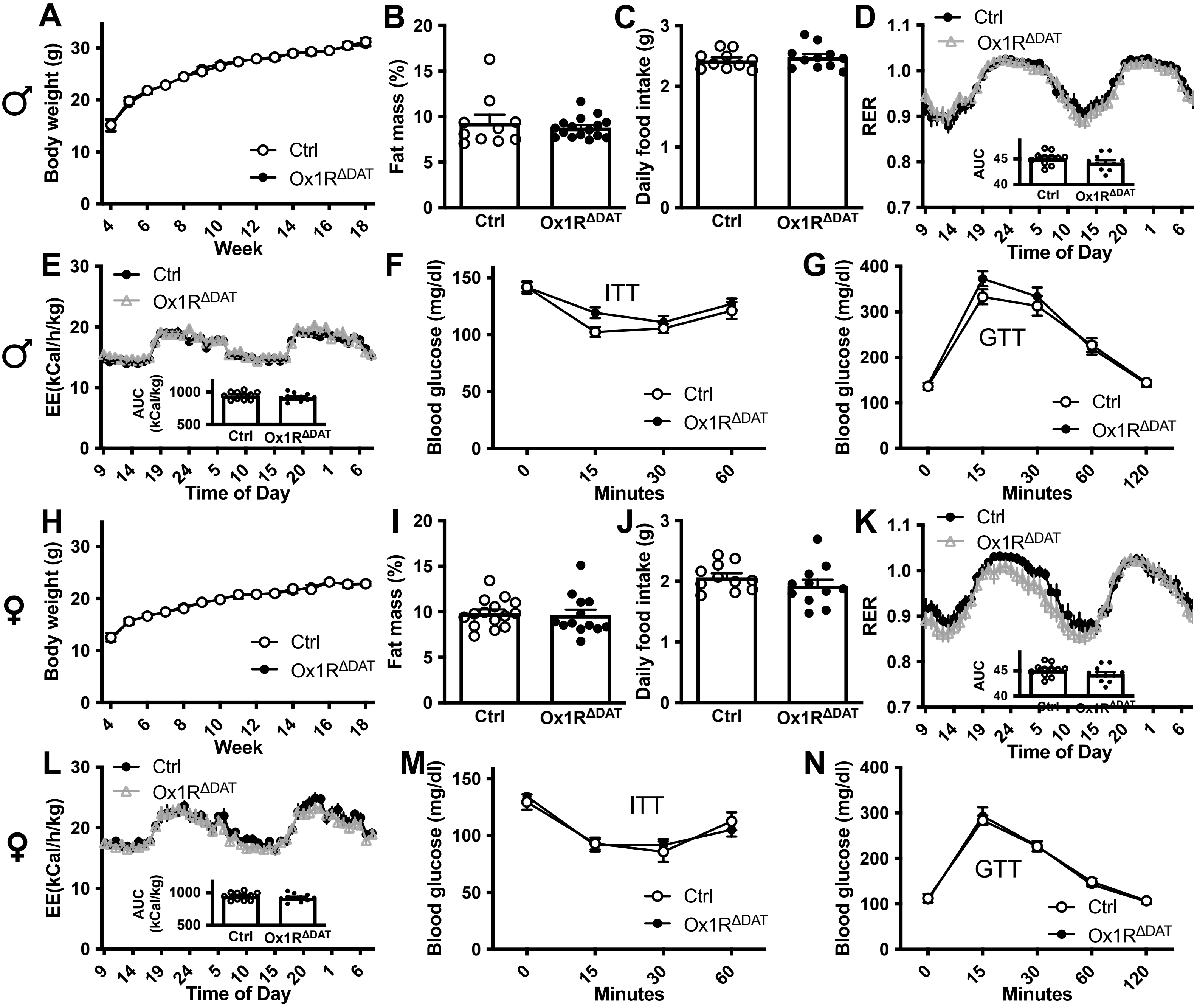


**Unaltered energy homeostasis in Ox1R$^{\Delta DAT}$ mice**.

**(A)** Average body weight (BW) of male mice. Control, n = 10; Ox1R$^{\Delta DAT}$, n = 12. **(B)** Average body fat content (percentage in BW) of male mice measured by nuclear magnetic resonance. Control, n = 10; Ox1R$^{\Delta DAT}$, n = 16. Phenomaster (PM) analysis of **(C)** daily food intake, **(D)** respiratory exchange ratio (RER) and **(E)** energy expenditure (EE) of male mice. Control, n = 10; Ox1R$^{\Delta DAT}$, n = 11 (C) or 12 (D, E). **(F)** Insulin tolerance test (ITT) of male mice. Control, n = 11; Ox1R$^{\Delta DAT}$, n = 12. **(G)** Glucose tolerance test (GTT) of male mice. Control, n = 16; Ox1R$^{\Delta DAT}$, n = 15. **(H)** Average BW of female mice. Control, n = 11; Ox1R$^{\Delta DAT}$, n = 11. **(I)** Average body fat content (percentage in BW) of female mice. Control, n = 15; Ox1R$^{\Delta DAT}$, n = 13. PM analysis of **(J)** daily food intake, **(K)** RER and **(L)** EE of female mice. Control, n = 11; Ox1R$^{\Delta DAT}$, n = 11. **(M)** ITT of female mice. Control, n = 10; Ox1R$^{\Delta DAT}$, n = 9. **(N)** GTT of female mice. Control, n = 10; Ox1R$^{\Delta DAT}$, n = 9. Data are represented as means ± SEM. The inserts in D, E, K and L present the area under curve (AUC) for statistical analysis. No significant differences were found between control and Ox1R$^{\Delta DAT}$ groups by two-way ANOVA (A, F, G, H, M, N) or unpaired two-tailed Student's tests (B-E, I-L). (A) Time: $F (14, 291) = 153.8$, $p < 0.0001$; genotype: $F (1, 291) = 0.13$, $p = 0.72$; time x genotype: $F (14, 291) = 0.21$, $p > 0.99$; (F) time: $F (3, 84) = 17.81$, $p < 0.0001$; genotype: $F (1, 84) = 3.70$, $p = 0.06$; time x genotype: $F (3, 84) = 0.97$, $p = 0.41$; (G) time: $F (4, 145) = 98.95$, $p < 0.0001$; genotype: $F (1, 145) = 1.38$, $p = 0.24$; time x genotype: $F (4, 145) = 0.94$, $p = 0.44$; (H) Time: $F (14, 247) = 102.4$, $p < 0.0001$; genotype: $F (1, 247) = 0.003$, $p = 0.96$; time x genotype: $F (14, 247) = 0.20$, $p > 0.99$; (M) time: $F (3, 68) = 18.57$, $p < 0.0001$; genotype: $F (1, 68) = 0.01$, $p = 0.92$; time x genotype: $F (3, 68) = 0.45$, $p = 0.72$; (N) time: $F (4, 85) = 131.7$, $p < 0.0001$; genotype: $F (1, 85) = 0.002$, $p = 0.96$; time x genotype: $F (4, 85) = 0.20$, $p = 0.94$. The full statistics are provided alongside the source data.

### Figure 3–figure supplement 1

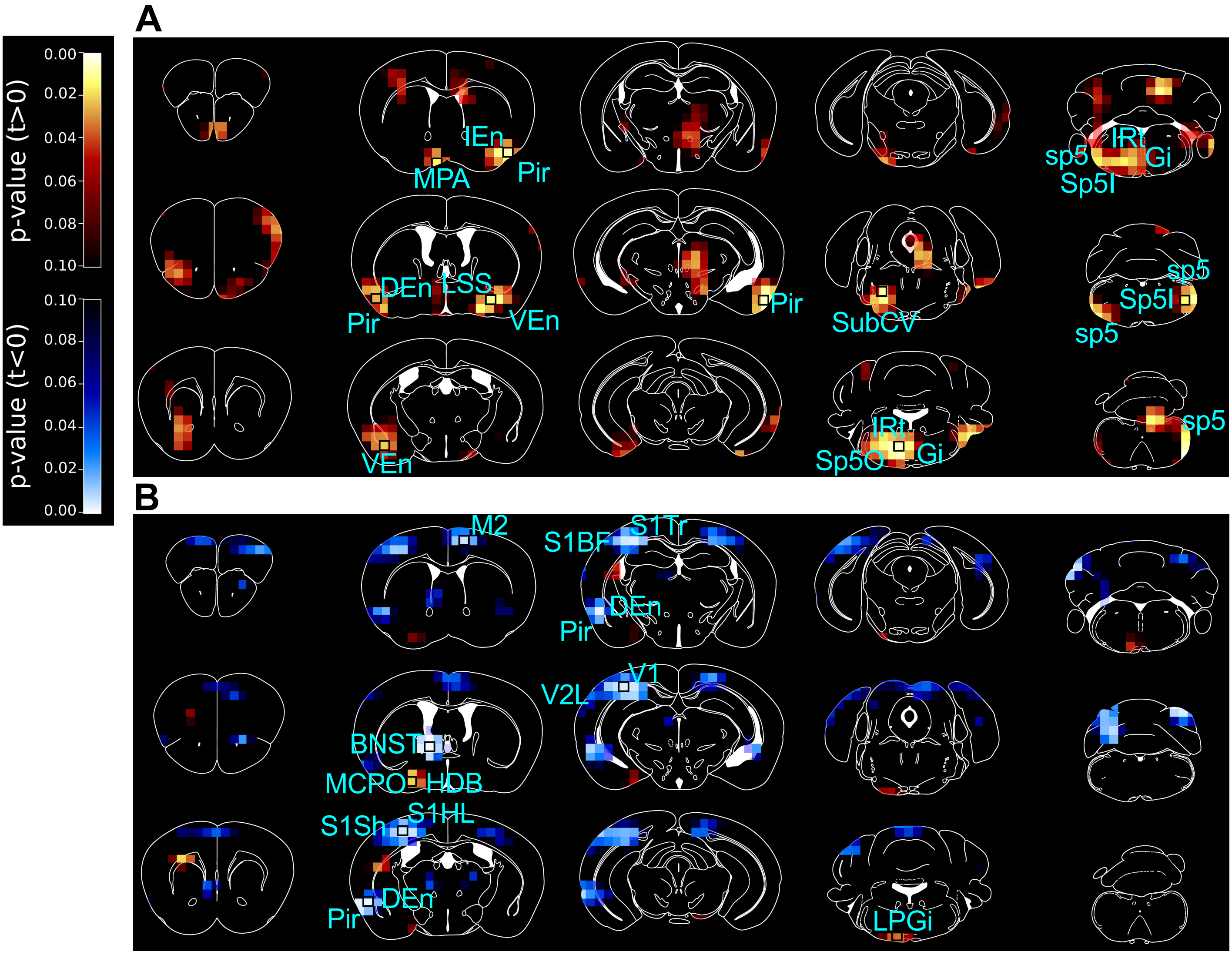


**PET imaging studies comparing Ox1R$^{\Delta DAT}$ and control mice, shown as p-map images with the original voxel size.**

3D maps of p-values in PET imaging studies comparing Ox1R$^{\Delta DAT}$ and control mice, after intracerebroventricular (ICV) injection of **(A)** saline (NS) and **(B)** orexin A. Brain areas with significant changes are indicated. Inserted black squares mark the voxels taken as examples to represent the respective area in Figure 3—supplement 2, which shows bar graphs with individual dots. Control-NS, n = 8; control-orexin, n = 6; Ox1R$^{\Delta DAT}$ (NS and orexin), n = 8. Data are compared by unpaired two-tailed Student's t-test. M2, secondary motor cortex; MPA, medial preoptic area; Pir, piriform cortex; IEn, intermediate endopiriform claustrum; DEn, dorsal endopiriform claustrum; VEn, ventral endopiriform claustrum; LSS, lateral stripe of the striatum; BNST, the dorsal bed nucleus of the stria terminalis; HDB, nucleus of the horizontal limb of the diagonal band; MCPO, magnocellular preoptic nucleus; S1Sh, primary somatosensory cortex, shoulder region; S1HL, primary somatosensory cortex, hindlimb region; S1BF, primary somatosensory cortex, barrel field; S1Tr, primary somatosensory cortex, trunk region; V1, primary visual cortex; V2L, secondary visual cortex, lateral area; SubCV, subcoeruleus nucleus, ventral part; Gi, gigantocellular reticular nucleus; IRt, intermediate reticular nucleus; LPGi, lateral paragigantocellular nucleus; Sp5O, spinal trigeminal nucleus, oral part; Sp5I, spinal trigeminal nucleus, interpolar part; sp5, spinal trigeminal tract.

### Figure 3–figure supplement 2

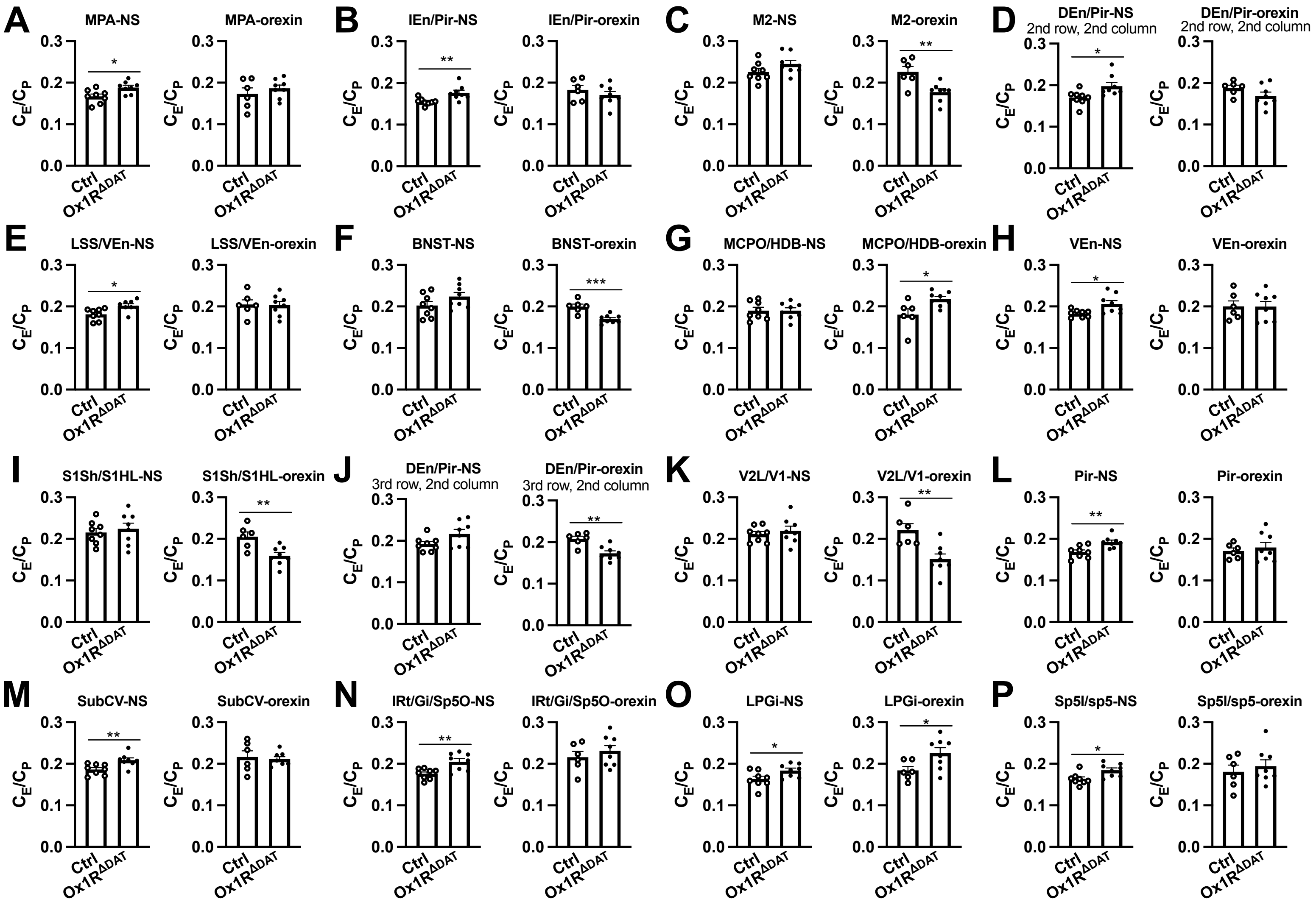


**Statistical testing for the volumes of interest from brain regions in PET imaging studies.**

The cerebral glucose ($C_E/C_P$) uptake, as measures for neuronal activity, is compared between $Ox1R^{\Delta DAT}$ and control (Ctrl) mice, after intracerebroventricular (ICV) injection of saline (NS) or orexin. The volumes of interest are from the brain regions labeled in the respective coronal planes of brain in Figure 3 and Figure 3–figure supplement 1. **(A)** MPA (medial preoptic area); **(B)** IEn (intermediate endopiriform claustrum) and Pir (piriform cortex); **(C)** M2 (secondary motor cortex); **(D)** DEn (dorsal endopiriform claustrum) and Pir, in the coronal planes shown in the 2nd row x 2nd column in Figure 3A and B and Figure 3–figure supplement 1A and B; **(E)** LSS (lateral stripe of the striatum) and VEn (ventral endopiriform claustrum); **(F)** BNST (the dorsal bed nucleus of the stria terminalis); **(G)** MCPO (magnocellular preoptic nucleus) and HDB (nucleus of the horizontal limb of the diagonal band); **(H)** VEn; **(I)** S1Sh (primary somatosensory cortex, shoulder region) and S1HL (primary somatosensory cortex, hindlimb region); **(J)** DEn and Pir, in the coronal planes shown in the 3rd row x 2nd column in Figure 3A and B and Figure 3–figure supplement 1A and B; **(K)** V2L (secondary visual cortex, lateral area) and V1 (primary visual cortex); **(L)** Pir; **(M)** SubCV (subcoeruleus nucleus, ventral part); **(N)** IRt (intermediate reticular nucleus), Gi (gigantocellular reticular nucleus) and Sp5O (spinal trigeminal nucleus, oral part); **(O)** LPGi (lateral paragigantocellular nucleus); **(P)** Sp5I (spinal trigeminal nucleus, interpolar part) and sp5 (spinal trigeminal tract). The row and column location described above the bar figures (D and J) indicates the respective coronal plane in Figure 3A and B and Figure 3–figure supplement 1A and B. Ctrl-NS, n = 8; Ctrl-orexin, n = 6; $Ox1R^{\Delta DAT}$, n = 8. Data are represented as means ± SEM. *$p < 0.05$; **$p < 0.01$; ***$p < 0.001$; as determined by unpaired two-tailed Student's t-test. The full statistics are provided alongside the source data.

### Figure 3–figure supplement 3

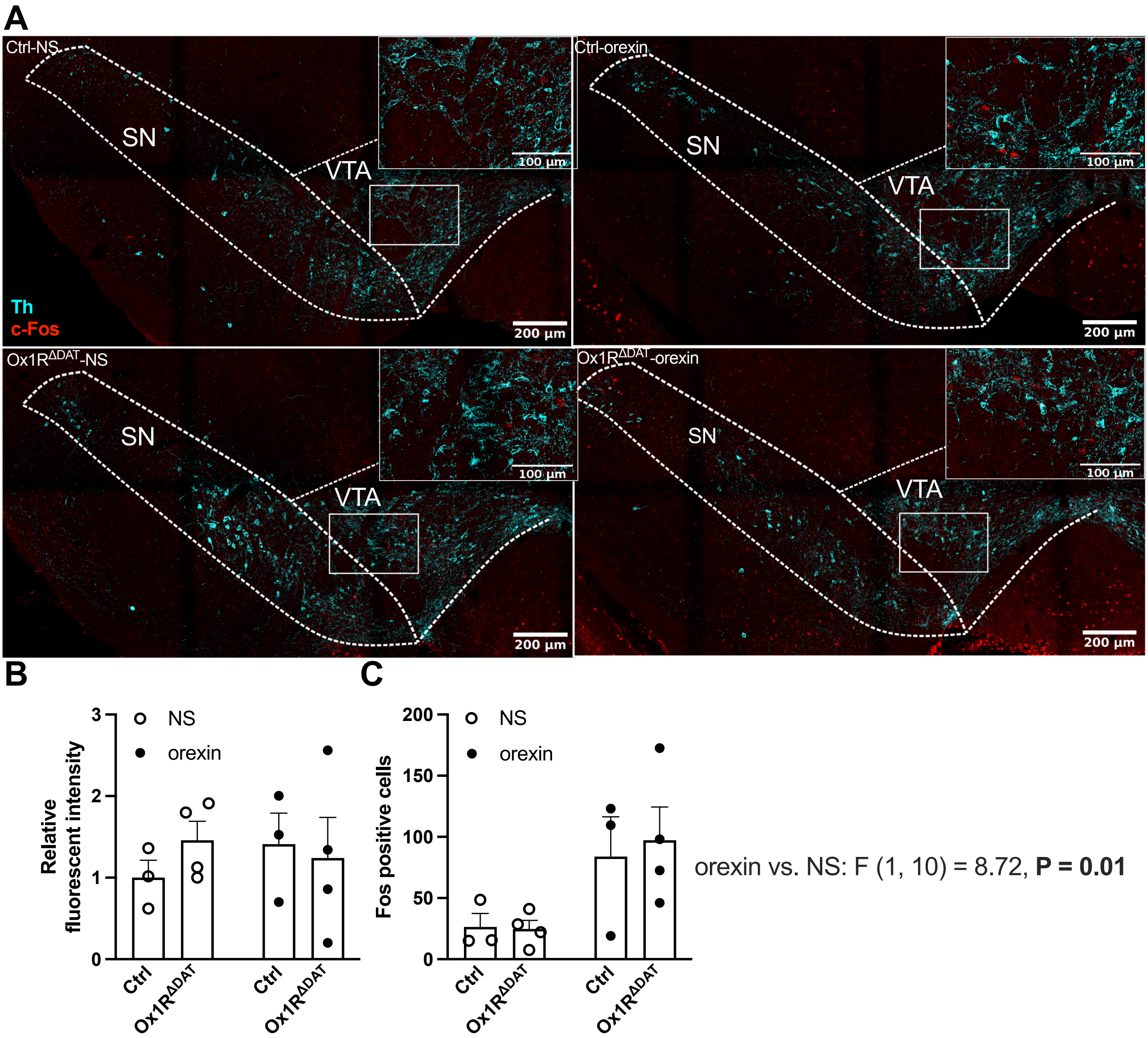


**c-Fos and tyrosine hydroxylase (Th) staining in substantia nigra (SN) and ventral tegmental area (VTA).**

**(A)** Representative images of c-Fos and Th staining in male control and Ox1R$^{\Delta DAT}$ mice injected (ICV) with saline (NS) or orexin A. **(B)** Quantification of Th fluorescence in SN and VTA. **(C)** c-Fos positive non-dopaminergic neurons in SN and VTA. Inserted images at the right up corner of each image are the amplifications of the marked area. Cyan, Th; red, c-Fos. Scale bar: 200 µm or 100 µm (insertions). Control, n = 3; Ox1R$^{\Delta DAT}$, n = 4. Data are represented as means ± SEM. p value is determined by two-way ANOVA followed by Sidak's post hoc test. (B) Treatment: F (1, 10) = 0.07, p = 0.80; genotype: F (1, 10) = 0.15, p = 0.71; treatment x genotype: F (1, 10) = 0.71, p = 0.42; (C) treatment: F (1, 10) = 8.72, p = 0.01; genotype: F (1, 10) = 0.07, p = 0.79; treatment x genotype: F (1, 10) = 0.12, p = 0.74. The full statistics are provided alongside the source data.

### Figure 4–figure supplement 1

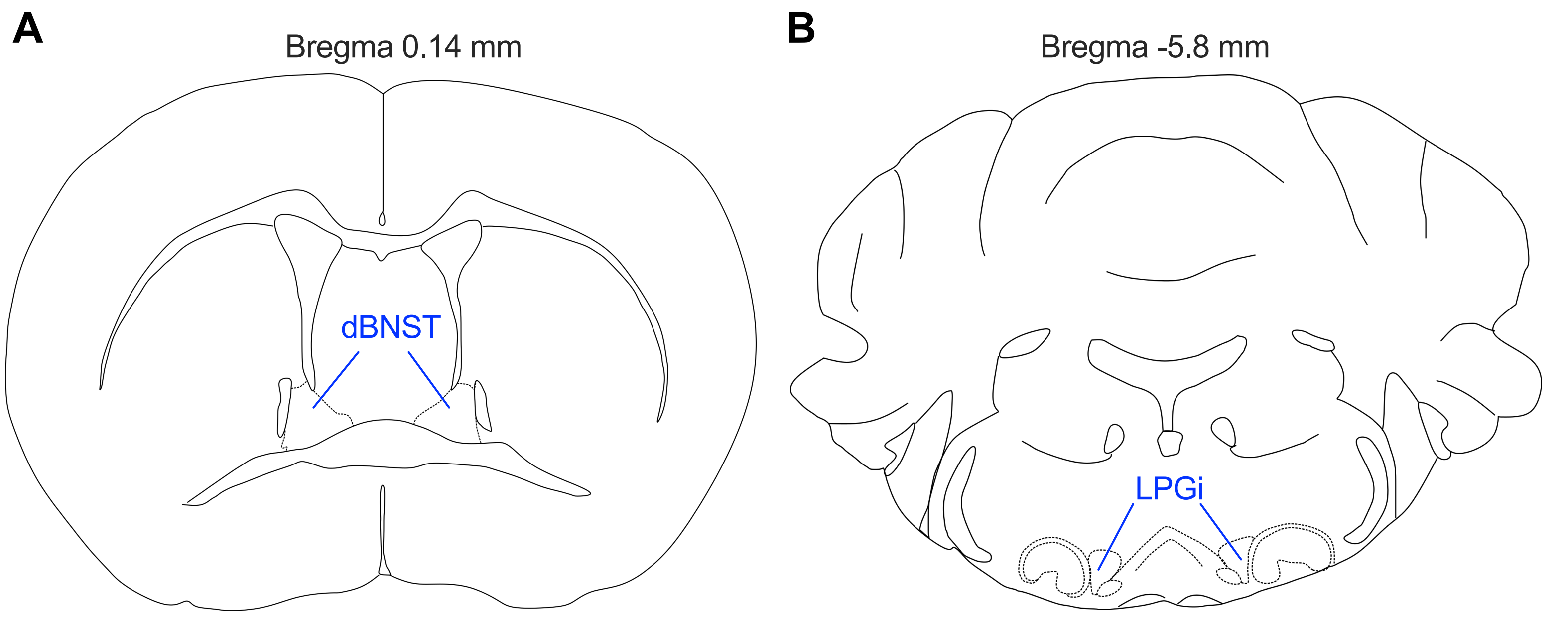


**A schematic illustration of the dorsal bed nucleus of the stria terminalis (dBNST) and lateral paragigantocellular nucleus (LPGi) based on the reference atlas.**

Blue text and lines indicate **(A)** dBNST and **(A)** LPGi. Unilateral nuclei were analysed and shown in Figure 4 and its supplements.

### Figure 4–figure supplement 2

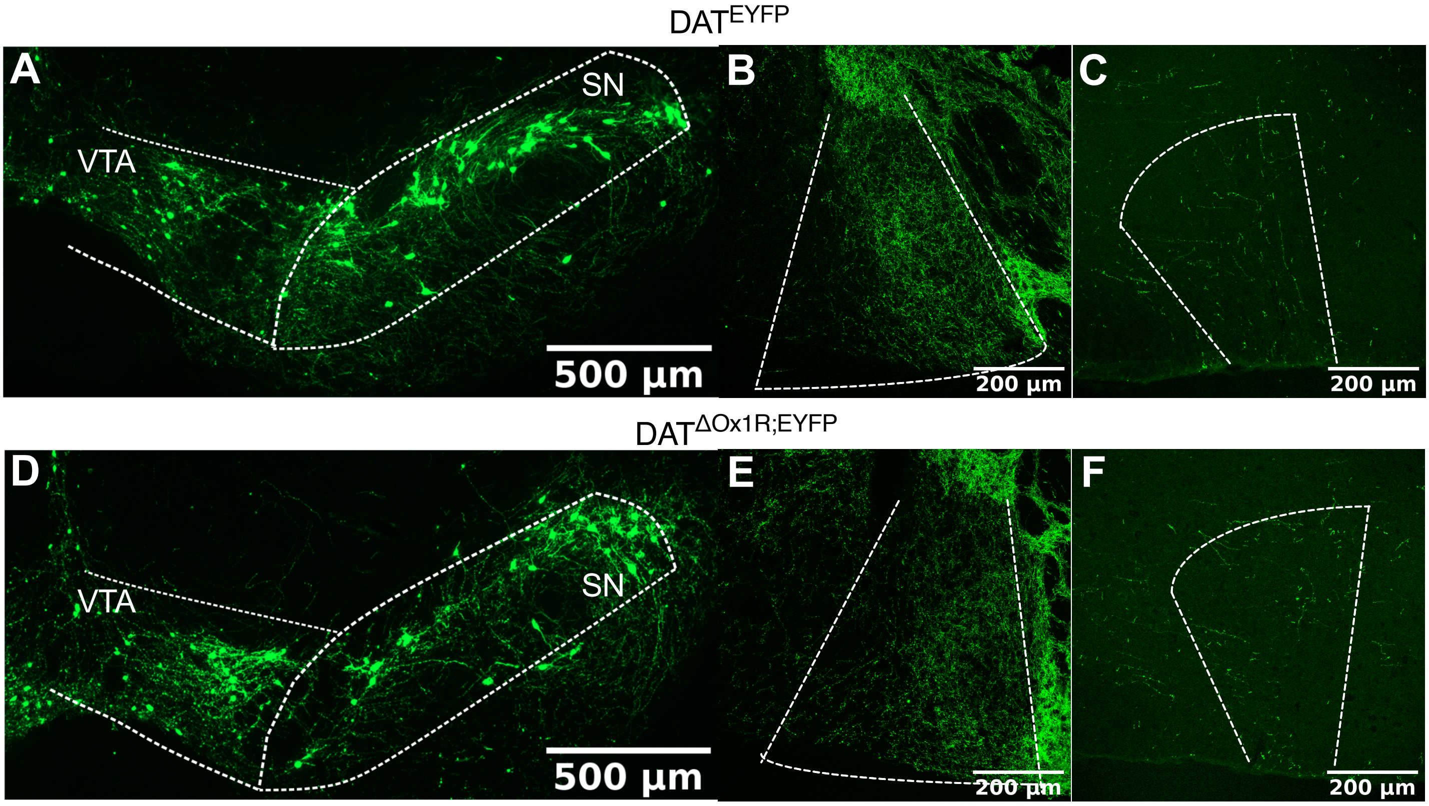


**Dopaminergic fibers innervating the dorsal bed nucleus of the stria terminalis (BNST) and lateral paragigantocellular nucleus (LPGi).**

Representative images of GFP staining in **(A)** substantia nigra (SN) and ventral tegmental area (VTA), **(B)** the dorsal BNST and **(C)** LPGi of $DAT^{EYFP}$ mice. Representative images of GFP staining in **(D)** SN and VTA, **(E)** the dorsal BNST and **(F)** LPGi of $DAT^{\Delta Ox1R;EYFP}$ mice. n = 3. Scale bar: 500 μm (A, D) or 200 μm (B, C, E, F).

### Figure 4–figure supplement 3

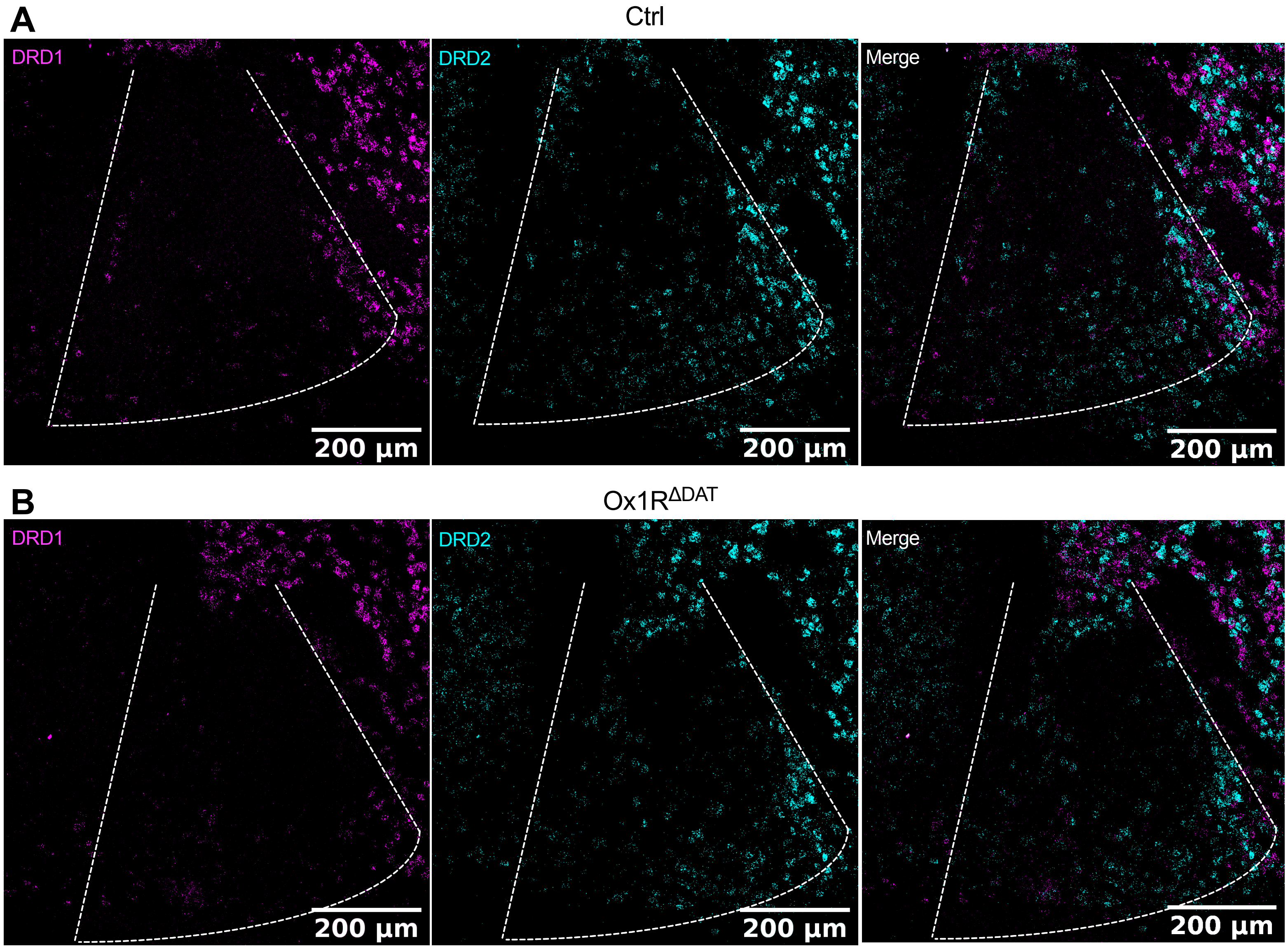


**Representative images showing dopamine receptors in the dorsal bed nucleus of the stria terminalis (BNST).**
**(A)** Representative images of D1 and D2 subtypes of dopamine receptor (DRD1 and DRD2) in the dorsal BNST of a control (Ctrl) mice. **(B)** Representative images of D1 and D2 subtypes of dopamine receptor (DRD1 and DRD2) in the dorsal BNST of an Ox1R$^{\Delta DAT}$ mice. Scale bar: 200 µm. Control, n = 3; Ox1R$^{\Delta DAT}$, n = 4. Magenta, DRD1; cyan, DRD2.

### Supplementary table 1

#### Functions of brain regions.

| Brain regions | Functions | References |
|---|---|---|
| piriform cortex, endopiriform claustrum, nucleus of the horizontal limb of the diagonal band, magnocellular preoptic nucleus | olfactory sensation | Smith et al., 2019; Wilson & Sullivan, 2011; Nunez-Parra et al., 2013 |
| spinal trigeminal nucleus and spinal trigeminal tract | pain and temperature sensation of oral cavity and the surface of face | Erzurumlu et al., 2010; Patel et al., 2022 |
| intermediate reticular zone | mastication | Nakamura et al., 2017; Travers et al., 2005 |
| gigantocellular reticular nucleus | motion related to arousal and motivation | Lemieux & Bretzner, 2019; Martin et al., 2011 |
| ventral part of subcoeruleus nucleus | sleeping | Hogl et al., 2018 |
| bed nucleus of the stria terminalis | emotion | Giardino et al., 2018 |
| lateral paragigantocellular nucleus | exploratory behaviours, locomotor speed | Takatoh et al., 2013; Arber & Costa, 2022; Capelli et al., 2017 |
| secondary motor cortex | action selection | Olson et al., 2020; Sul et al., 2011 |
